# Structural basis of CO_2_ valence coding in *Drosophila*

**DOI:** 10.64898/2026.01.05.697655

**Authors:** Dominik Javorski, Beate Bergkirchner, Gernot Ensinger, Alexander Lingl, Jelena Navolic, Ashwaq Batawi, Thomas Hummel

**Affiliations:** Department of Neurosciences and Development, University of Vienna, Djerassiplatz 1, 1030 Vienna, Austria; Center for Molecular Neurobiology Hamburg, University Medical Center Hamburg-Eppendorf, Falkenried 94, 20251 Hamburg, Germany; King Abdulaziz University, 21589 Jeddah, Saudi Arabia

## Abstract

In the olfactory system, glomerular sensory channels of single receptor identity support reliable odor recognition for appropriate approach or avoidance behaviors. For many olfactory stimuli, the assigned sensory value is innate but weighted by the internal state and previous experiences. How context-dependent modulation of innate valence coding supports distinct behavioral responses is poorly understood. Here we show that CO_2_ sensory information in *Drosophila*, intrinsically aversive but modified by attractive food signals, diverges from the canonical glomerular channel already in the antennal lobe via the polarized local interneuron LN23. LN23 relays sensory input via an extraglomerular CO_2_ pathway and manipulation of LN23 activity revealed a dominant role in CO_2_-induced avoidance behavior. The extraglomerular CO_2_ pathway projects to the posterior lateral protocerebrum (PLP) adjacent to the canonical Lateral Horn (LH) olfactory processing center and segregates into anatomically distinct valence channels. Connectome data together with functional characterization identified convergence of parallel CO_2_ channels onto interconnected third-order neurons that are anatomically positioned to receive information from additional sensory modalities via distinct pathways: while the glomerular CO_2_ pathway converges with food relay neurons onto separated dendritic domains of the PD5 interneuron in the LH, the extraglomerular pathways with anatomically identified connections to CO_2_-associated and humidity-/temperature-related circuits are positioned to provide potentially antagonistic inputs onto the PLP interneuron PV9. This early anatomical divergence of a defined olfactory channel and subsequent convergence in higher brain centers provide a structural framework for investigating context-dependent valence coding and behavioral flexibility.

## Introduction

The ability to recognize and react to environmental stimuli is essential for the survival of animals. Insect olfactory systems allow the sensing and integration of volatile signals to identify food resources and avoid potential threats (Wilson, 2013). In insects, local changes in atmospheric CO_2_ concentration can induce context-dependent behavioral responses (Faucher et al. 2006; Hansson & Stensmyr, 2011). Walking *Drosophila* show robust CO_2_ induced avoidance (Suh et al. 2004; Kwon et al. 2007; Faucher et al. 2006), while in more aroused states such as flight or foraging, CO_2_ becomes an attractive olfactory cue (van Breugel et al. 2018) which is linked to dopamine levels affecting mushroom body output neurons (Lewis et al. 2015; Sachse & Beshel 2016). At the circuit level, prolonged CO_2_ exposure can reduce avoidance behavior; previous studies have implicated plasticity of GABAergic interneurons in olfactory habituation (Das et al. 2011; Olsen & Wilson, 2008; Nagel & Wilson, 2016; Sachse et al. 2007; Semelidou et al. 2018). Interestingly, *Drosophila* avoidance behavior directly correlates with the activity of CO_2_ specific olfactory receptor neurons (MacWilliam et al. 2018), raising the question of how CO_2_-induced sensory activity is transformed into central valence representations and modulated to support context-dependent behavior (Frechter et al. 2019).

The peripheral CO_2_ circuit follows the canonical organization of olfactory sensory channels. A single population of ORNs (ab1C) co-express the gustatory receptors Gr21a and Gr63a and projects ipsilaterally to the V glomerulus in the ventral antennal lobe (Suh et al. 2004; Kwon et al. 2007; Vosshall and Stocker 2007). CO_2_ sensation is transmitted to higher brain centers through two glomerular PNv types: a bilateral cholinergic PN (PNv^bi^) projecting to both the Mushroom Body (MB) and Lateral Horn (LH), and a LH specific relay via a unilateral PN (PNv^uni^) (Grabe et al., 2016; Lin et al. 2013; Sachse et al. 2007; Tanaka et al. 2012). Although distinct spatial LH domains encoding opposite hedonic values and odor intensity have been identified (Frechter et al. 2019; Das Chakraborty et al. 2022), glomerular PNs provide synaptic input to a diverse set of LH neurons making the identification of central brain representation and context-dependent modulation of a given sensory channel difficult (Dolan et al. 2019; Scheffer et al. 2020).

Here we describe the identification of a CO_2_ relay channel, morphologically and functionally separate from the canonical glomerular circuit, via a polarized olfactory interneuron, LN23. The LN23 pathway mediates a robust CO_2_-induced aversion largely unaffected by glomerular circuit plasticity and provides an anatomical route for possible early multimodal interactions. Furthermore, the parallel pathways of glomerular and extraglomerular PNs converge onto a small group of LH interneurons as a core domain for CO_2_ valence coding in the central brain.

## Material and Methods

### Drosophila Stocks

VT031497 (PNv^bi^ Gal4 and the alternative line: R53A05; BDSC#38859; BDSC: Bloomington *Drosophila* Stock Center), NP7273 (PNv^uni^ Gal4), R44A02 (LN23 Gal4; BDSC#50196) were used to target neurons PNv^bi^, PNv^uni^ and LN23 respectively. LN23-lexA (BDSC#53644) and PNv^bi^-lexA (VT031497 lexA) were used in sybGRASP (BDSC#93197 / reverse direction BDSC#64315) and tGRASP (BDSC#79040/reverse direction BDSC#79039) experiments to express the postsynaptic GFP fragment. Gr21-Gal4 (BDSC#57600) was used to target ORNs which project specifically to the V-glomerulus. To target LNv1 and LNv2, we used NP1227P from the Max Planck Institute and NP2426-P from the Kyoto Stock Center. The Gal4 driver line BDSC #40444 was used for l2LN19. UAS-shibirets (Gift from Barry Dickson) was used to inhibit neurons at restrictive 31°C. UAS-ReaCh (BDSC#53741) was used in optogenetic experiments to artificially activate specific neuronal subsets which are labelled by the Gal4 lines. CantonS and w1118 wildtype flies were used for control experiments. The Gr63a1 (BDSC#9941) mutant stock was used in the behavior experiments as control for flies which are unable to detect CO_2_. In addition to our main LN23 line (BDSC#50196) an additional line also labeling LN23 (BDSC#48408) was used for optogenetics. UAS-CaLexA (BDSC#66542) was used in combination with the Gal4 lines to label active neurons. To further activate or silence specifically active neurons a CaLexA stock without GFP was used (BDSC#66543) in combination with lexAop-ReaCh or lexAop-shits respectively. UAS-DenMark::cherry was used to label dendrites of neurons while UAS-syb::GFP and UAS-syt::GFP specifically labeled presynaptic terminals. To confirm the location of active T-bars in LN23 and PNv neurons, we also applied UAS-brp::GFP (BDSC#36291 and BDSC#36292).

### Immunohistochemistry

Flies were dissected in PBS (phosphate buffer solution) and brains were fixed in 2% PFA (Paraformaldehyde). After the fixation brains were washed 3 times with 0.3% PBT (phosphate buffer with TritonX) and then blocked with Goat serum. After 1 hour of blocking the 1st AB (Antibody) was applied and the brains were kept in a 4°C fridge overnight. The next day brains were washed again 3 times before applying the 2nd AB. After the staining protocol was complete, brains were mounted and afterwards scanned with a confocal microscope. Anti Ncad rat (1:10, DHSB) was used to label all neuropils in the fly brain in all experiments. For CaLexA experiments anti GFP rabbit (1:5000, Invitrogen) was used additionally to enhance the signal strength, according to Masuyama et al. (2012). GRASP experiments, DenMark and labeling of presynaptic terminals (syt::GFP and syb::GFP) didn”t require any antibodies besides anti-Ncad since the endogenous signal was sufficient and changes after CO_2_ incubation should not be distorted. For MCFO, first Antibodies were Anti-Ncad rat (1:10) or Anti-ollas rat (1:2000), Anti-flag mouse (1:2000) and Anti-HA rabbit (1:2000). As second antibodies anti-rat 647 (1:500), anti-mouse (highly cross-absorbed, 1:300) and anti-rabbit (1:500) were applied.

### MCFO (Multicolor flip-out)

To label single cells in a usually rather broad expressing Gal4-line, we took advantage of a method established by Nern et al. 2015, called MCFO. They recombined four constructs, consisting of an UAS-site, followed by a stop-cassette to suppress transcription and one of four different transmembrane proteins, which can, if expressed, be labeled via antibodies. These four independent constructs can also be expressed together in the same cell, which leads to a mixture of colors and increased probability of one cell getting labeled in one distinct color. The stochastic excision of the stop cassette is achieved by a heat sensitive enzyme, whose activation is triggered by incubating the flies at 37°C in a water bath.

### GRASP

To visualize synaptic contacts, two different GRASP tools were used. SybGRASP is activity dependent because one fragment of GFP is located on vesicle membranes. As vesicles fuse with the presynaptic membrane, which is dependent on incoming action potentials, the large GFP fragment is exposed in the synaptic cleft and can reconstitute with the small GFP fragment located on the postsynaptic membrane (Macpherson et al. 2015). tGRASP, or targeted GRASP (Shearin et al. 2018), is activity independent and the two GFP fragments are bound to pre and postsynaptic specific proteins (Cacophony and Telencephalin respectively). Comparison of these two GRASP techniques provided complementary indicators of activity-associated contacts and anatomical apposition, but neither assay directly measured synaptic currents or the number of functional synapses. No antibodies against GFP were used since the endogenous expression was sufficient. GFP intensities were quantified with FIJI, which is measured as the mean gray value of a defined region of interest (ROI) above a threshold. To subtract the mean grey value of the background noise, intensity was measured in another glomerulus (same size of ROI) which was not showing any signal. Each datapoint in the diagrams represents one quantified glomerulus. The same applies to intensity measurements for the following two sections.

### DenMark::cherry / Syt:GFP, syb::GFP

UAS-syt::GFP and UAS-syb::GFP were used to label presynaptic terminals of target neurons. Synaptotagmin (syt) and synaptobrevin (syb) are proteins of the vesicle membrane and therefore a Gal4 driven expression of the GFP tagged proteins shows strong signal in the presynaptic terminals. UAS-DenMark (Dendritic Marker) is a somatodendritic marker (cherry) to label dendrites. GFP/Cherry intensities were quantified with FIJI.

### CaLexA

CaLexA (Masuyama et al. 2012) was used to label active cells. It is based on the NFAT transcription factor, which is calcium responsive. As Ca+ enters a cell, a chimeric lexA-VP16-NFAT, enters the nucleus where expressed lexA can bind to lexAop inducing the expression of GFP. Stronger or more prolonged calcium-dependent activation is expected to increase accumulated GFP, although reporter intensity cannot resolve brief activity dynamics or be equated directly with firing rate. GFP levels were quantified in FIJI as it was described in the section about GRASP. There was no selection for gender and all flies tested were 2 days old at the start of the experiments. For chronic CO_2_ exposure experiments, flies were maintained at 5% CO_2_ for three days for CaLexA activity mapping, whereas structural measurements of V-glomerulus volume and ORNv axonal morphology were performed following five days of exposure. This was because we wanted to follow established experimental paradigms for CO_2_-induced V-glomerulus plasticity, particularly those described by Sachse and colleagues (2007). We also used a CaLexA construct where expressed lexA is not expressing GFP but instead driving the expression of a temperature sensitive shibire protein to silence neurons. Further we also used this stock in combination with lexA driven expression of ReaCh to specifically further activate physiologically active neurons. 99% spermidine was used as an inhibitor to GR21a receptors. 5 µL were put on a small piece of cotton which was then transferred to a vial containing the flies which were used in the experiments after 3 days of exposure. For the brief-stimulation condition shown in Figure 1O, flies were exposed to 100% CO_2_ for 30 seconds on the laboratory microscope CO_2_ pads. After the flies recovered from anesthesia, the exposure was repeated twice more, for three 30-second exposures in total; the interval between pulses was determined by recovery rather than a fixed time. Ambient air refers to laboratory air containing approximately 0.04% CO_2_, whereas chronic exposure refers to the experimental 5% CO_2_ condition for three days unless otherwise specified.

**Figure 1.**
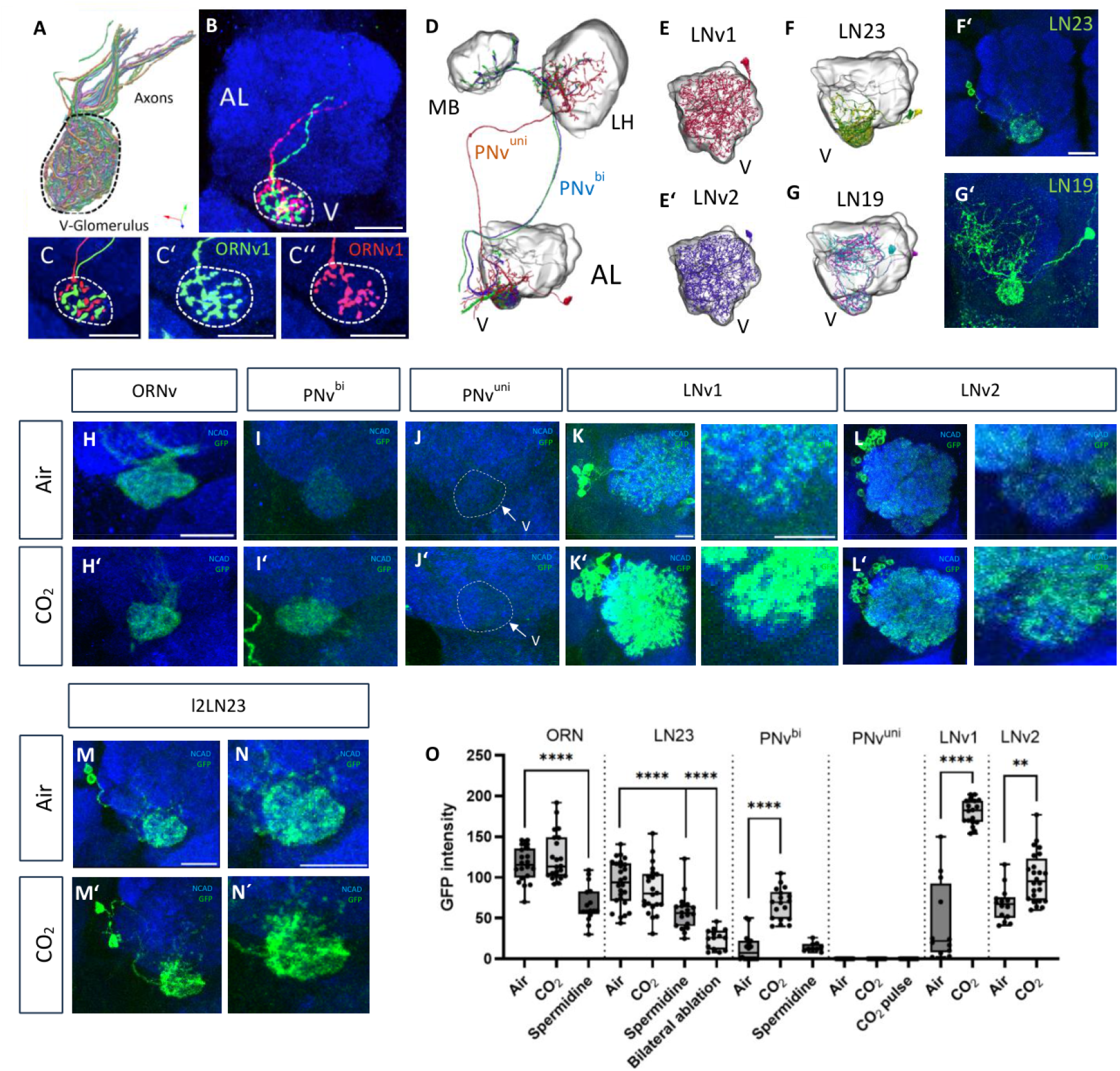
Principal neurons of the V glomerulus show distinct response properties for CO_2_. (A–C) About 40 olfactory sensory neurons (ORNv) innervate the ipsilateral V glomerulus at the ventral AL surface (A). Single-cell clones revealed the terminal branch pattern of ORNv axons (B). Each ORNv axon forms 3–5 non-overlapping terminal branches (C). (D–G) Morphology of central neurons of the V glomerulus, including two types of projection neurons (PNv^uni^ and PNv^bi^, D), two classes of broad local interneurons (LNv1 and LNv2, E/E⍰), and two classes of sparse LNs (LN23 and LN19, F, G). Gal4 expression lines for sparse interneurons LN23 and LN19 (F⍰, G⍰). (H–N) CaLexA reporter expression in different neuron classes under ambient air (H–N) and elevated CO_2_ levels (H⍰–N⍰). Under ambient air conditions, CaLexA labeling was detected in ORNv, LNv1/2, and LN23, but little or no signal was detected in the two glomerular PN classes. (H⍰–N⍰) Following chronic CO_2_ stimulation, ORNv and LN23 retained strong CaLexA-GFP labeling, comparable to their signal under ambient-air conditions. Based on CaLexA-GFP intensity, reporter labeling increased in PNv^bi^, LNv1, and LNv2 following prolonged exposure, whereas PNv^uni^ showed no detectable CaLexA-GFP accumulation. (O) Quantification of GFP signal intensity in ORNv, LN23, PNv^bi^, PNv^uni^, LNv1, and LNv2. Flies were maintained in ambient air or exposed to 5% CO_2_ for three consecutive days. For ORNv, LN23, and PNvbi, spermidine was applied under ambient-air conditions to inhibit GR21a-mediated CO_2_ detection. Bilateral antennal ablation was included as an additional sensory-deprivation condition for LN23. For the pulsed condition tested in PNv^uni^, flies received three 30-second exposures to 100% CO_2_ on the laboratory microscope CO_2_ pads, with recovery to wakefulness between successive exposures. The absence of GFP accumulation in this condition does not exclude a rapid neuronal response. Scale bars: 20 µm, unless otherwise indicated.

### Behavioral experiments

For behavior experiments, flies were kept in total darkness, in 25°C incubators in vials containing fly food. Flies which were used in optogenetics were raised on fly food containing all-trans-retinal to enhance their response to light exposure (Vilinsky et al. 2018). All flies were age matched and tested when they were 3-4 days old. Since there was no observed difference between male and female flies both sexes were used in all behavioral experiments. As we did not observe any behavioral artifacts caused by either green or red LED light we opted for the green LEDs, since they elicited a stronger response in the flies (compared to Inagaki et al., 2014). Food was present during rearing, but food odor was not independently manipulated as an experimental stimulus in the CO_2_/optogenetic choice assay.

Flies were immediately transferred to empty vials which can be connected to the CO_2_ or optogenetic setup after they were taken out of the incubators. The whole procedure was also done in darkness (red light was used to be able to handle the flies, which is barely visible for the flies) to minimize visual distraction affecting the fly”s behavior. The whole apparatus was shifted sideways, and the flies were gently moved to the elevator segment of the setup. The elevator was lowered to a position short above the decision point and the flies were given 60 seconds rest. After the resting period either a green LED light or CO_2_ flow was activated and the elevator was lowered to the decision point. For CO_2_ preference experiments, the test arm was supplied with 5% CO_2_, whereas the control arm received ambient air.

The animals were given 20 seconds to move inside the behavioral setup and make a choice. Afterwards the elevator was pulled up again, so that the flies remained in either the test tube, control tube or the central elevator area. Each of these populations was counted and a preference index (PI) calculated, that was defined as the percentage of flies located in the control arm (air filled /dark) of the setup, minus the percentage of flies located in the test arm (CO_2_ filled/light). Since in some of the tested populations, a few animals remained in the central elevator part of the arena, their decision was defined as neutral. Their number was divided by two and added to both the test and control population for the PI. Each datapoint in the diagrams represents the PI of a group of flies (n=15-40). Assigning neutral flies equally to the two arms leaves the PI numerator unchanged and retains all tested flies in the denominator; thus PI = 100 × (Ncontrol − Ntest)/Ntotal. Therefore, we split the non-choosing flies equally between both arms because they did not demonstrate a preference. This ensures that they do not bias the index toward either side, while still accounting for the fact that not every fly made a choice.

### Hemibrain analyses

Connectivity data were obtained from the neuPrint interface (neuPrint, Janelia Research Campus; https://neuprint.janelia.org/) built on the Hemibrain connectome (Scheffer et al. 2020). It was used to study and model the neurons involved in the V-circuit. The main goal was to identify synaptic connectivity between the different layers of information processing within the circuit. Finally, the putative circuit was reconstructed in 3D and PN/LN output zones were identified. Neuronbridge.janelia.org was used to gather 3D images from l2_PNm17 and sm_PNm1. FlyWire.ai was a tool to quickly get information about the predicted neurotransmitter identities of neurons.

### Statistics

All diagrams and statistical analyses were done by using GraphPad Prism 9 software. Normally distributed data were compared with a student”s t-test, whereas datasets that did not meet normality assumptions were analyzed with a Mann–Whitney U test. Behavioral data spanning multiple days were tested with a one-way ANOVA. Statistical significance is indicated in the figures by asterisks, with p < 0.05 (*), p < 0.01 (**), p < 0.001 (***), and p < 0.0001 (****).

## Results

### Glomerular circuit dynamics of CO_2_ perception

The V glomerulus at the ventral AL surface is innervated by a single population of about 40 CO_2_-sensitive olfactory receptor neurons (ORNv) expressing the two gustatory receptors GR21a and GR63a (Kwon et al. 2007; Jones et al. 2007) (Fig.1A-B). Each ORNv axon forms 3-5 terminal branches tightly packed in a non-overlapping fashion with other ORNv terminals (Fig. 1, C-C’’), providing sensory input to glomerular inter- and projection neurons (Fig1D-G’) (Grabe et al., 2016). To identify relay neurons of the V glomerulus with different physiological properties we screened a collection of Gal4 lines using the CaLexA activity reporter to compare accumulated calcium-dependent reporter labeling under ambient-air conditions (Koyama et al. 2021; Masuyama et al. 2012) (Fig. 1H-O). Gal4 lines for ORNv show strong CaLexA reporter activity (Fig.1H), consistent with previous evidence that CO_2_-receptive sensory neurons differ in presynaptic inhibition (MacWilliam et al. 2018; Olsen & Wilson, 2008; Root et al. 2008). In contrast to the strong reporter labeling in ORNv, the two uniglomerular PN classes of the V glomerulus, unilateral V_l2PNs (PNv^uni^) and bilateral V_ilPNs (PNv^bi^, Fig. 1D) do not show any significant level of CaLexA activation under ambient air conditions (Fig. 1I, J). Two main classes of local interneurons, LNv1 and LNv2 display moderate levels of CaLexA activity at atmospheric CO_2_ levels (Fig. 1K, L). A strong CaLexA-GFP signal, comparable to ORNv can be observed for l2LN23 (LN23), a less characterized olfactory class of sparsely innervating local interneurons (Fig.1F, F”, M and N) (Liou et al. 2018). Fluorescence signal of Ca2+-induced GFP expression quickly decreases in LN23 following sensory deprivation in adult flies via removal of both antennae (Fig.1O). Furthermore, an extended exposure to the ORNv inhibitor spermidine (MacWilliam et al., 2018) results in a reduced CaLexA activity not only in ORNv, but also LN23, consistent with an association between antennal sensory input and LN23 reporter activation under ambient-air conditions (Fig. 1O).

A predominant relay of CO_2_-induced sensory activity by LN23 is supported by the modified activity-dependent GRASP technique (Macpherson et al. 2015). Expression of the presynaptic syb::GFP1-10 in ORNv and the postsynaptic GFP11 in AL neurons revealed a significantly stronger GRASP signal for ORNv→LN23 connectivity compared to the ORNv→PNv^bi^ synaptic activity (Fig.2F and G). Similarly, the activity-independent tGRASP (Shearin et al. 2018) is most prominent for ORNv→LN23 (Fig.2I and J), which is in line with connectome data (neuPrintExplorer, hemibrain; Scheffer et al. 2020; Schlegel et al. 2021). Together, these reporter measurements identify LN23 as a prominent ORNv-associated circuit component, with a stronger GRASP signal than that observed for PNv^bi^ under the tested conditions.

### Cell type plasticity of glomerular relay neurons following chronic CO_2_ stimulation

We next determined how the overall activity of V glomerulus relay- and interneurons changes following long-lasting sensory stimulation. Adult *Drosophila melanogaster* avoids an experimentally presented CO_2_ stimulus relative to an ambient-air control in a T-maze choice assay (Faucher et al. 2006; Suh et al. 2004) (Fig. 2A) but shows reduced avoidance following chronic 5% CO_2_ exposure (Das et al. 2011, Fig. 2B, C). The reduction of CO_2_-induced aversive responses is associated with an increase in the synaptic volume of the V glomerulus (Fig.2D; Sachse et al. 2007 and Das et al. 2011). At the ORN level, mosaic analysis revealed an extension in the surface area and size of individual synaptic boutons of ORNv axons (Fig.2E). Compared to ambient CO_2_ levels, prolonged elevated CO_2_ (5%) sensation results in neuron-type specific changes of CaLexA activity (Fig. 1 H”-N”) (Masuyama et al. 2012). While PNv^bi^ neurons show a strong increase in Ca2+ levels following prolonged CO_2_ exposure (Fig.1I’), no CaLexA-dependent GFP expression could be observed in PNv^uni^ (Fig.1J’), indicating distinct CaLexA reporter profiles for the two main glomerular CO_2_ relay channels (Lin et al. 2013). In addition to chronic CO_2_ stimulation, three 30-second exposures to 100% CO_2_ also did not produce detectable CaLexA-GFP expression in PNv^uni^ (Fig.1O). No significant changes in CaLexA-GFP levels could be detected for LN23 neurons following prolonged CO_2_ exposure compared to the already high expression at ambient CO_2_ conditions (Fig1M’, N’, O).

**Figure 2.**
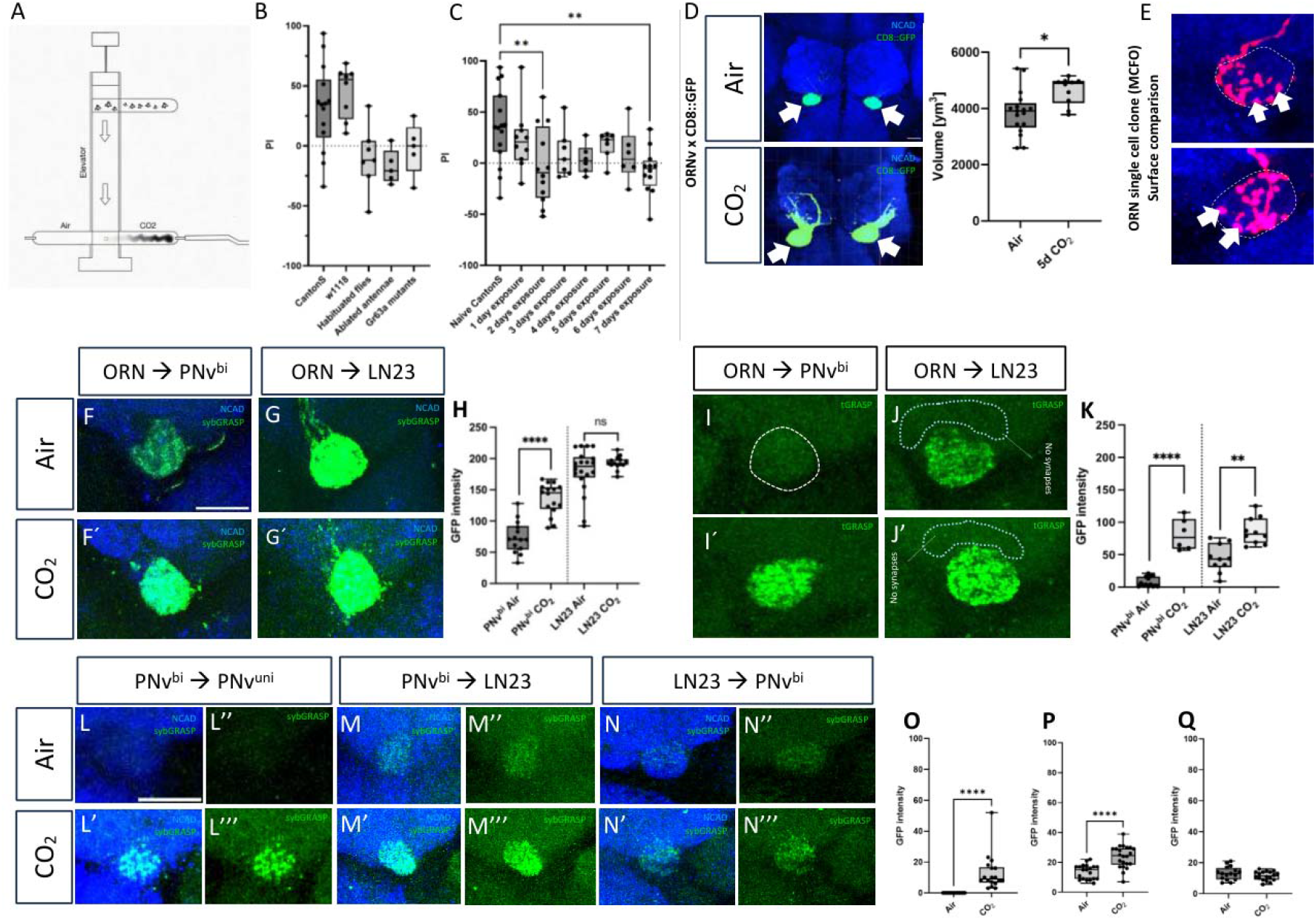
Circuit dynamics of the V glomerulus following prolonged CO_2_ exposure. (A) T-maze setup to determine CO_2_ preference in adult flies. (B) Preference index (PI) of naïve flies compared with flies subjected to chronic CO_2_ exposure, bilateral antennal removal, or mutation of the CO_2_ receptor GR63a; each condition reduced innate avoidance of the test CO_2_ stimulus. A value of +100 indicates that all flies selected the air arm, whereas −100 indicates that all flies selected the 5% CO_2_ arm. Flies remaining in the central elevator were counted as neutral and allocated equally to both arms when calculating PI, as described in Methods. (C) Change in CO_2_ avoidance during prolonged exposure; avoidance was already reduced after one day. (D, E) Structural changes in V-glomerulus volume (D) and ORNv axonal arborization (E) following the five-day exposure used for these structural measurements. (F–Q) Synaptic connectivity among neurons of the V glomerulus analyzed by transgenic GRASP. Activity-dependent sybGRASP indicates enhanced ORNv→PNv^bi^ connectivity (F⍰, H), but no significant changes for ORNv→LN23 (G⍰, H) following prolonged exposure. Activity-independent tGRASP expression indicates increased reporter labeling for ORNv connections with PNv^bi^ and LN23, consistent with morphological changes in ORN terminals (I⍰, J⍰, K). ORN input is restricted to the glomerulus and was not detected in the extraglomerular region (arrows, J). Panels L and O quantify PNv^bi^ →PNv^uni^ sybGRASP, panels M and P quantify PNv^bi^ →LN23 sybGRASP, and panels N and Q quantify LN23→PNv^bi^ sybGRASP. The first two comparisons increased following prolonged exposure, whereas the LN23→PNv^bi^ comparison showed no significant change.

**Figure 3.**
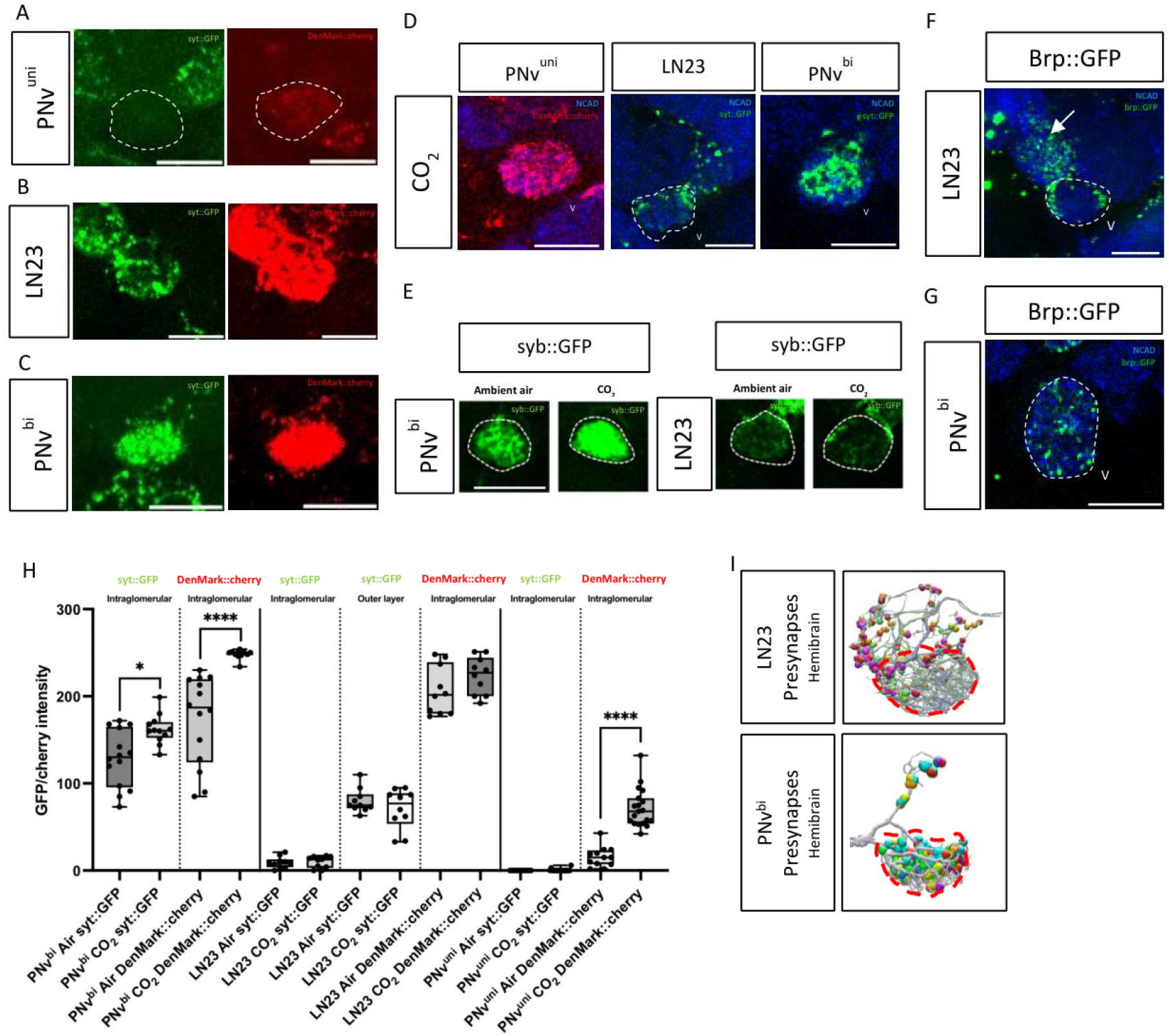
Neuronal polarity of CO_2_ glomerular circuit components. (A–C) Confocal sections show syt::GFP and DenMark::mCherry expression in PNv^uni^, LN23, and PNv^bi^. Quantification of GFP/mCherry levels revealed distinct localization of presynaptic sites in PNv^bi^, PNv^uni^, and LN23. PNv^bi^ displayed prominent presynaptic labeling throughout the glomerular neuropil, while PNv^uni^ showed no detectable presynaptic signal despite clear dendritic arborizations. LN23 exhibited scattered presynaptic puncta largely outside the glomerular boundaries. (D) PNv^uni^, LN23, and PNv^bi^ show different responses following prolonged CO_2_ exposure. LN23 did not show increased syt::GFP or DenMark::mCherry signal, while both increased for PNv^bi^; for PNv^uni^, only DenMark::mCherry increased. (E) Syb::GFP labeling further showed that presynaptic sites in PNv^bi^ were concentrated in the glomerular neuropil and increased after chronic CO_2_ exposure, whereas LN23 signals remained scattered along the glomerular margin. (F, G) Higher-resolution optical sections show brp::GFP puncta in LN23 processes outside the V glomerulus (F, arrow), but confined to the V glomerulus in PNv^bi^ (G). (H) Quantification of syt::GFP and DenMark::mCherry. For LN23, intensity measurements at the center of the glomerulus and near the glomerular boundary were distinguished. (I) Hemibrain reconstructions show the enrichment of LN23 presynaptic sites (colored spheres) in extraglomerular domains, compared with their intraglomerular distribution in PNv^bi^.

As GABAergic local interneurons are associated with olfactory habituation (Das et al. 2011; Sadanandappa et al. 2013) we compared changes in CaLexA activation in LNv1 and LNv2 (Das et al. 2011, Sachse et al. 2007). While an increase in the Ca2+ reporter signal could be observed for both LN types following chronic CO_2_ exposure, the CaLexA reporter signal of LNv1 is significantly higher than that of LNv2 (Fig.1K” and L’). In line with the cell-type specific CaLexA expression, previous connectome analysis showed significant differences of LN input for CO_2_ relay neurons: (neuPrintExplorer, hemibrain): While both PNv^uni^ and LN23 do not receive major LN-type input, PNv^bi^ receives anatomically identified input from putatively GABAergic lLN1 neurons (“bc” subtype, 16% of input) and a group of multiple lLN2 neuron types (X, F, P, 19%) (Schlegel et al. 2021; Bates et al. 2020). These results indicate cell type specific physiological changes of glomerular neurons following prolonged CO_2_ exposure, in which PNv^bi^ shows dynamic changes in glomerular connectivity while both PNv^uni^ and LN23 neurons showed no comparable increase in CaLexA labeling relative to ambient-air controls. We next tested how the cell-type specific changes in neuronal activity after prolonged exposure alter their GRASP signals within the V glomerulus. A strong increase of activity-dependent sybGRASP levels for ORNv input onto PNv^bi^ could be detected following chronic CO_2_ stimulation (Fig.2F”). In contrast, no significant changes in sybGRASP levels were observed for ORNv input onto LN23 (Fig.2G’). Of note, compared to ambient-air conditions, the standard deviation for ORNv→LN23 sybGRASP levels was reduced after prolonged CO_2_ exposure, although the mean reporter signal did not increase significantly (Fig.2H). In addition, activity-independent tGRASP expression increased at contacts between ORNv and the two main relay neurons, PNv and LN23 (Fig.2I”, J”), which is in line with morphological changes of ORN axon terminals described above (Fig.2D, E). Following prolonged CO_2_ exposure, the PNv^bi^ class displayed increased sybGRASP levels on PNv^uni^ and LN23 (Fig.2L-M”““), while no significant changes could be observed for LN23 on PNv^bi^ (Fig.2N-N”““). Taken together, CO_2_-induced ORNv activity seems to be relayed via two distinct glomerular output channels dependent on the stimulus strength: LN23 and PNv showed distinct reporter responses across the tested exposure conditions.

### Identity, innervation patterns and polarity of 2nd order CO_2_ sensitive neurons

To test if the difference in CaLexA activity might be related to the organization of presynaptic sites within CO_2_ relay neurons we visualized the active zones using syt::GFP/brp::GFP transgenes together with the dendrite marker DenMark::mCherry (Kittel et al. 2006, Nicolai et al. 2010; Wagh et al. 2006). PNv^bi^, PNv^uni^ and LN23 significantly differ in their expression and localization of presynaptic proteins within the dendritic compartment (Fig.3A-C). While PNv^bi^ shows prominent dendritic syt::GFP localization throughout the glomerulus (Fig.3C), no syt::GFP signal can be detected within PNv^uni^ dendrites (Fig.3A), indicating different cellular polarity of the two main glomerular PNv classes. Despite the significant expression level, syt::GFP appeared more scattered within LN23 dendrites and less restricted to the glomerulus neuropil compared to PNv^bi^ (Fig.3B). In fact, single optical sections allow the localization of the active zone marker Brp::GFP in LN23 dendrites outside the glomerular neuropil (Fig.3F) but in PNv^bi^ confined to the V glomerulus (Fig.3G). In line with these results hemibrain EM data lists enriched presynaptic site in LN23 dendrites outside the V glomerulus in contrast to the restriction PNv^bi^ presynaptic sites to the glomerular neuropil (Fig.3I) (https://neuprint.janelia.org/; Schlegel et al. 2021; Bates et al. 2020).

Following chronic sensory stimulation, PNv^bi^ showed a significant increase in the density of presynaptic sites while no significant change in the amount presynaptic syt::GFP labeling of LN23 dendrites can be observed (Fig.3D). Like the subcellular organization at ambient CO_2_ conditions, presynaptic sites of PNv^bi^ dendrites are uniformly distributed throughout the V-glomerulus (Fig.3D, last image), while most presynaptic sites of LN23 dendrites are found outside of the glomerular boundaries (Fig.3D, middle image). In contrast to CO_2_ relay neurons PNv^bi^ and LN23, no dendritic localization of presynaptic marker transgenes could be detected for PNv^uni^ (Fig.3D, first image), neither under ambient nor chronic CO_2_ condition, further supporting a differential synaptic profile of the two PN classes. All syt::GFP and DenMark::cherry intensities were quantified using ImageJ (Fig.3H). An additional presynaptic marker syb::GFP revealed a similar expression pattern: while PNv^bi^ showed a strong signal within the glomerular boundaries, which increases following CO_2_ incubation, LN23 dendrites show a robust GFP signal adjacent to the neuropil border of the V glomerulus (Fig.3E). As both, PNv^bi^ and PNv^uni^ are described as being excitatory, cholinergic neurons (Bates et al. 2020; Lin et al. 2013; Schlegel et al. 2021; Tanaka et al. 2012) we investigated the neurotransmitter identity of LN23. Anti-GABA immunostaining does not label LN23 cell bodies (Supplemental Fig.3A). This does not rule out other inhibitory mechanisms (see discussion). These results revealed not only main differences in the neuronal organization among the 3 classes of CO_2_ relay neurons, but also a distinct response profile in intraglomerular plasticity of dendro-dendritic connections. CO_2_ input diverges into two more robust feed-forward channels (PNv^uni^ towards LH and LN23 within the AL) while the dynamic PNv^bi^ pattern indicates intraglomerular processing.

### Developmental profile of the inter-glomerular CO_2_ relay neuron

Having established the polarity of LN23, our next objective was to investigate the development of this neuron, specifically focusing on how interglomerular relay neurons are integrated into stereotyped antennal lobe circuits. While the glomerulus-specific assembly of olfactory receptor neuron (ORN) axons with projection neuron (PN) dendrites has been extensively documented (Hong & Luo 2014), less is known about the mechanisms underlying the incorporation of relay neurons into these circuits. At the beginning of pupal development, PN dendrites innervate the early precursor of the adult AL and occupy distinct spatial domains before ORN axons enter the AL by around 20h APF (Jefferis et al. 2001; Jefferis et al. 2004). LN23 neurons can be traced back to the late larval stage with a cellular morphology related to SEZ (subesophageal zone) interneurons (Truman & Bate, 1988; Winding et al. 2023, Fig. 4). During early pupal stages, larval LN23 precursor neurons retract from the SEZ to accumulate at the ventral region of the AL precursor neuropil (Fig4A-C”). Here LN23 processes target a region which is devoid of dendrites of uniglomerular PNs (Fig4F, F”). Before ORN axon arrival, LN23 neurons establish a separate dendritic branch, which extends dorsally before bending back to the posterior AL and merges with the main dendritic domain (Fig. 4D-E”).

**Figure 4.**
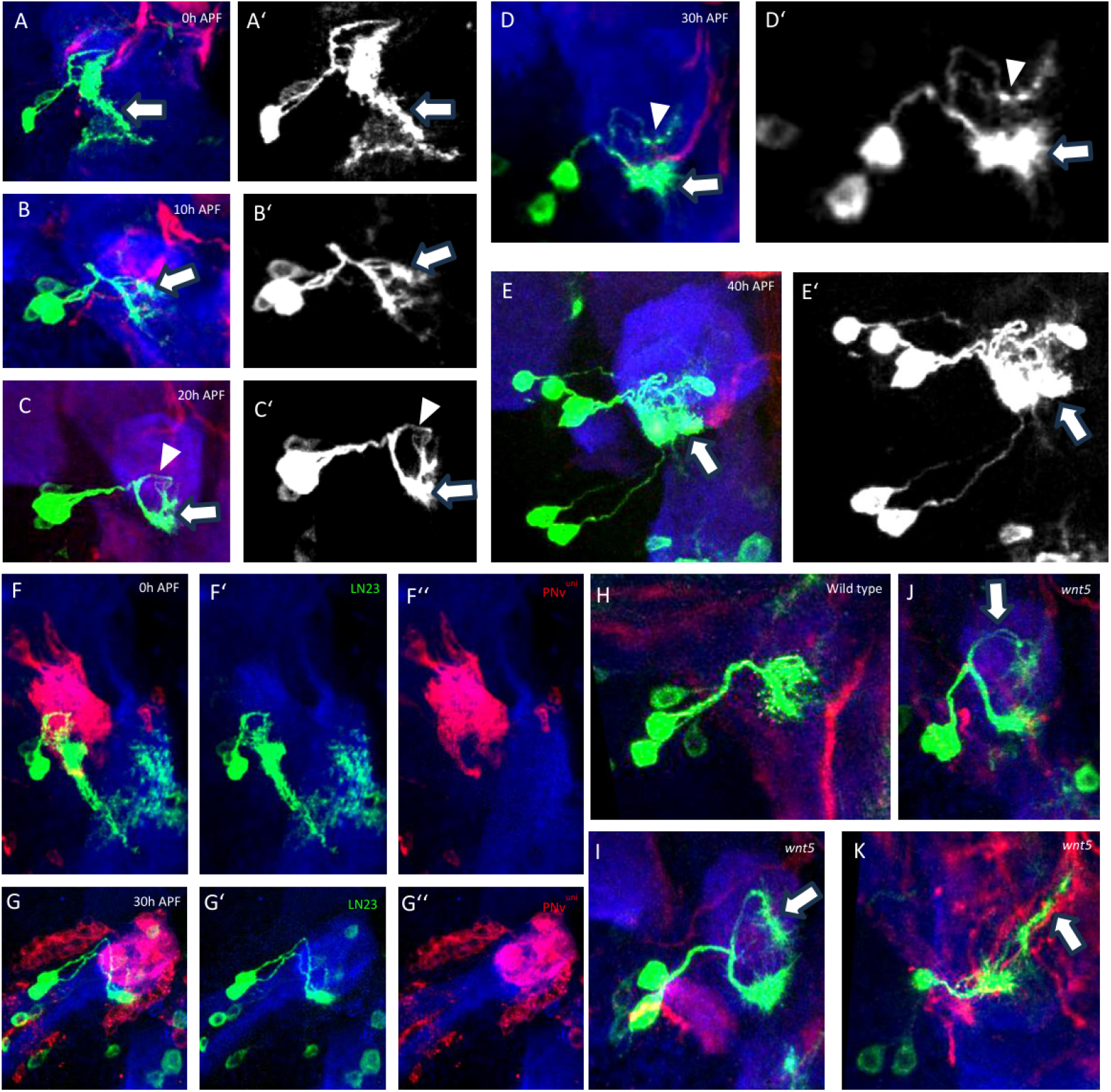
Early LN23 development. (A–E⍰) At the beginning of pupal development (0 h APF), larval LN23 neurons remodel their ventral processes (arrow in A) and start extending into the developing adult AL (arrow in B). While a broad arborization can be observed in the ventral AL (arrows in C and D), a small process extends into the dorsal AL to target a region posterior to the V glomerulus (arrowheads in C and D). At 40 h APF, the main LN23 dendrites are restricted to the glomerular boundaries (arrow in E). (F–G⍰ ⍰) During reorganization from the larval to the adult AL, LN23 processes (green) develop in proximity to uniglomerular PN dendrites (red). Compared with wild-type (H), wnt5 mutants (I–K) show greater separation of the ventral and dorsal LN23 branches (arrows in I and J), which subsequently extend along axon tracts toward higher brain centers (arrow in K).

Loss of the secreted molecule Wnt5, which affects PN dendrite patterning (Hing et al. 2020; Wu et al. 2014), leads to a continued growth of the dorsal LN23 branch along PN axon tracts which exit the AL towards higher olfactory brain centers (Fig.4I-K). These results indicate that LN23 neurons display an intrinsic polarity similar to uniglomerular PNs, but instead of relaying sensory information from AL glomeruli to the LH/MB, a Wnt5-dependent mechanism seems to prevent AL exit and target the presynaptic process to the extraglomerular region in the ventral AL. In wild type, ORN axons innervate the developing V glomerulus only at the ipsilateral AL, in contrast to the majority of bilateral ORN classes (Fig.4F-H) (Couto et al. 2005). To gain insight into the mechanisms which support glomerulus-specific ORNv input on LN23, we analyzed candidate genes which are expressed on growing ORNs. Mutations in the protocadherin-type cell adhesion molecule Flamingo (fmi), which is highly enriched on ORN axons in the developing V glomerulus, affect their synaptic targeting. Here, fmi mutant ORNv axons are no longer restricted to the glomerular boundaries but extend into the posterior AL next to the output domain of LN23 (Suppl.Fig.4A, B), indicating cellular interactions to restrict class-specific axon-dendrite recognition within the glomerular domain.

In summary, LN23 shows cellular and developmental features that resemble those of olfactory projection neurons: a highly polarized neuronal morphology in the adult olfactory system is established by a glomerulus-specific dendrite targeting for singular ORN input and a default axonal extension along olfactory PN tracts rerouted by external signals (Komiyama & Luo, 2006).

### LN23 mediates CO_2_-induced aversion

To test the role of the different CO_2_ relay neurons on *Drosophila* behavior, we expressed the shibirets transgene to block synaptic transmission in a cell type specific manner (Chen et al. 1991; Kitamoto 2001). Neuronal silencing of each of the two glomerular PNs led to a reduction of the innate CO_2_ avoidance, with stronger effects observed for PNv^uni^ compared to PNv^bi^ (Fig. 5A). While CO_2_ avoidance can still be observed following the neuronal silencing of each PN class, shibire^ts^-induced silencing of LN23 reduced CO_2_ avoidance to near-neutral levels under the tested conditions (Fig. 5A). To test for cell type specificity of the behavioral phenotype following LN23 silencing, we used the Ca2+ dependent LexA activation of the CaLexA construct for shits expression and observed a reduction of the innate CO_2_ aversion (Fig. 5B) (Masuyama et al. 2012). To increase lexA levels we chronically exposed flies to high levels of CO_2_. Interestingly, increased CaLexA mediated shits expression via chronic exposure not only suppressed behavioral aversion but led to a significant level of attraction towards CO_2_ (Fig.5B). These results indicate a prominent function of LN23 in CO_2_ valence coding: Blocking LN23 activity is sufficient to fully suppress CO_2_ behavioral aversion and following chronic CO_2_ exposure in the activity-dependent silencing condition, the PI shifted toward attraction. The contribution of PNv^bi^ cannot be separated from accompanying exposure-dependent changes in this comparison (Jones et al. 2007; Kwon et al. 2007, MacWilliam et al. 2018; Suh et al. 2004).

**Figure 5.**
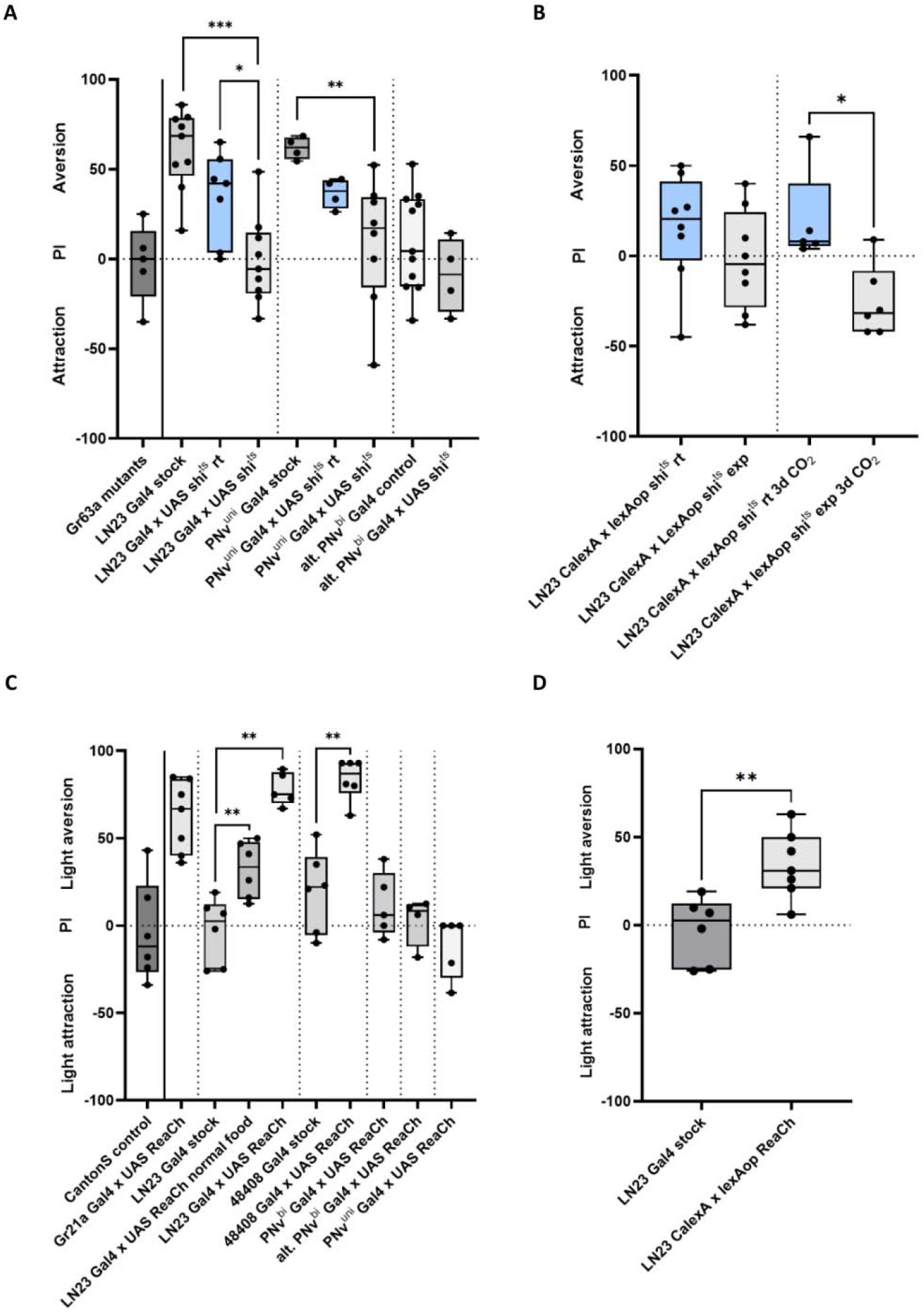
Activity of LN23 controls CO_2_ aversion behavior. (A) Cell type-specific silencing with UAS-shibirets revealed that neuronal activity of all three V-glomerulus relay neurons contributes to innate CO_2_ avoidance. Silencing of PNv^uni^ or PNv^bi^ reduced avoidance behavior, with a stronger effect in PNv^uni^. In contrast, LN23 silencing almost entirely abolished CO_2_ aversion under the tested conditions. Control experiments performed at room temperature (rt; exp denotes the restrictive-temperature condition), where shibirets should remain permissive, also showed some reduction in avoidance in PNv^bi^ and LN23 genotypes. Since experiments with PNv^uni^-Gal4 (VT031497) showed inconclusive results, the alternative PNv^bi^ driver R53A05 (BDSC#38859) was used. (B) CaLexA-dependent expression of shibirets in the LN23-targeted population reduced CO_2_ avoidance; following prolonged CO_2_ exposure, flies showed attraction rather than avoidance. (C) Optogenetic activation using UAS-ReaChR showed that LN23-targeted activation elicited aversive behavior in the choice assay. Light activation of ORNv (Gr21a-Gal4) and LN23 (independent LN23-Gal4 and 48408-Gal4 drivers) triggered avoidance, while activation of PNv^bi^ or PNv^uni^ failed to elicit significant changes. Flies were reared on food, but food odor was not independently manipulated as a stimulus in this assay. (D) CaLexA-driven ReaChR expression in LN23 also induced light-mediated aversion, providing a complementary activity-dependent manipulation.

To further determine if these relay neurons are sufficient for behavioral aversion, we optogenetically activated each neuron type and measured the behavioral response (Inagaki et al. 2014; Klapoetke et al. 2014; Pulver et al. 2009) (see M&M). Light-induced activation of ORNv (GR21a-Gal4) expressing UAS-ReaCh led to strong aversion (Fig. 5C). Similarly, optogenetically activated LN23 is sufficient to trigger behavioral aversion via two independent Gal4 driver lines (LN23 Gal4 and the alternative line we used: VT48408-Gal4; Fig.5C) or by Ca2+ induced LexA activation (Fig.5D). In contrast, no change in light-induced activation for both, PNv^bi^ (two independent driver lines, see Methods) nor PNv^uni^ could be observed (Fig.5C). These results support a predominant role of LN23 in CO_2_-induced avoidance behavior. Compared to the more dynamic activity of PNv^bi^ following acute and chronic sensory stimulation, LN23 shows little changes in intrinsic activity, with comparatively small changes in the integrated CaLexA reporter across the conditions tested. Based on these findings we propose a working model in which LN23 makes a major contribution to CO_2_-evoked aversion, while parallel PN channels may modulate the behavioral response.

### 3rd order neurons postsynaptic to LN23 have a putative role in valence coding

To gain further insights into the unique organization of the LN23 pathway compared to the glomerular relay channel, we analyzed the postsynaptic candidate neurons within the posterior AL. Two PN types, l2_PNm17 and sm_PNm1 are receiving a large proportion of their synaptic input (27% and 18%, respectively) from LN23 (Fig.6A, neuPrint, Janelia Research Campus; https://neuprint.janelia.org/; Schlegel et al. 2021). Like the two PN classes of the V glomerulus, PNm1 and PNm17 show distinct relay properties: While input into PNm1 is mostly restricted to the LN23-CO_2_ channel, dendrites of PNm17 extend into adjacent VP1 and VP3-5 glomeruli to receive direct input from thermo- and humidity-receptor neurons (neuPrint, Janelia Research Campus; https://neuprint.janelia.org/; Frank et al. 2015; Marin et al. 2020). Both types of non-olfactory relay neurons target multiple distinct brain areas adjacent to the canonical olfactory LH, the posterior lateral protocerebrum (PLP), superior clamp (SCL) and the superior lateral protocerebrum (SLP) (Fig6 A, B, C) (Ito et al. 2014). To visualize the trajectories of the PNs, we used MCFO (Aso et al. 2014) to create single cell clones, which show that PNm1 and PNm17 axons turn close before entering the LH to target adjacent brain regions (Fig. 6B, C). Of note, compared to glomerular PNv^uni^ and PNv^bi^, PNm1 and PNm17 receive only minor input from GABAergic LNs (neuPrint, Janelia Research Campus; https://neuprint.janelia.org/), indicating reduced gain control properties of the extra-glomerular ORN→LN23→PNm pathway (Olsen & Wilson, 2008).

**Figure 6.**
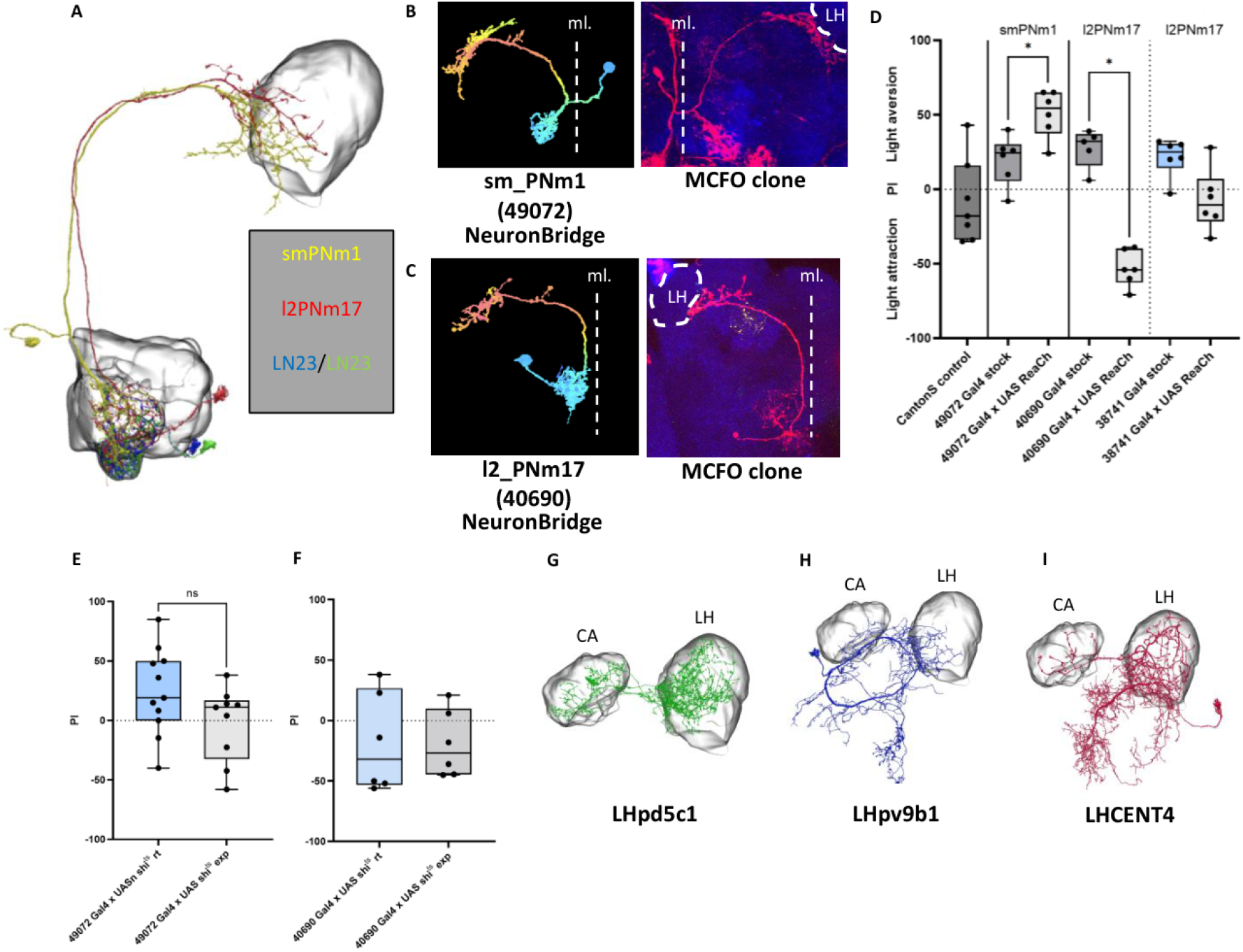
Two postsynaptic neurons of LN23 elicit opposing behavioral responses when activated. (A) Connectomic analysis (neuPrint, Janelia Research Campus) revealed two major postsynaptic partners of LN23, PNm1 and PNm17. (B, C) Single-cell MCFO clones of sm_PNm1 and l2_PNm17 show axonal projections bypassing the canonical olfactory lateral horn (LH) and targeting adjacent regions including the posterior lateral protocerebrum (PLP), superior clamp (SCL), and superior lateral protocerebrum (SLP). (D) Optogenetic activation of PNm1 induced strong aversion, whereas PNm17 activation elicited attraction, under the optogenetic choice-assay conditions. (E, F) Silencing experiments with shibirets attenuated CO_2_ avoidance after PNm1 silencing, whereas PNm17 silencing did not significantly alter the response. (G–I) Hemibrain connectome data identified distinct downstream LH partners for the two CO_2_ relay pathways: LHPD5c1 as a major postsynaptic target of PNv^bi^, LHPV9b1 downstream of PNm1, and LHCENT4 downstream of PNv^uni^. “rt” indicates room-temperature control, while “exp” indicates the restrictive-temperature condition. The midline is indicated by the dashed line (ml.).

**Figure 7.**
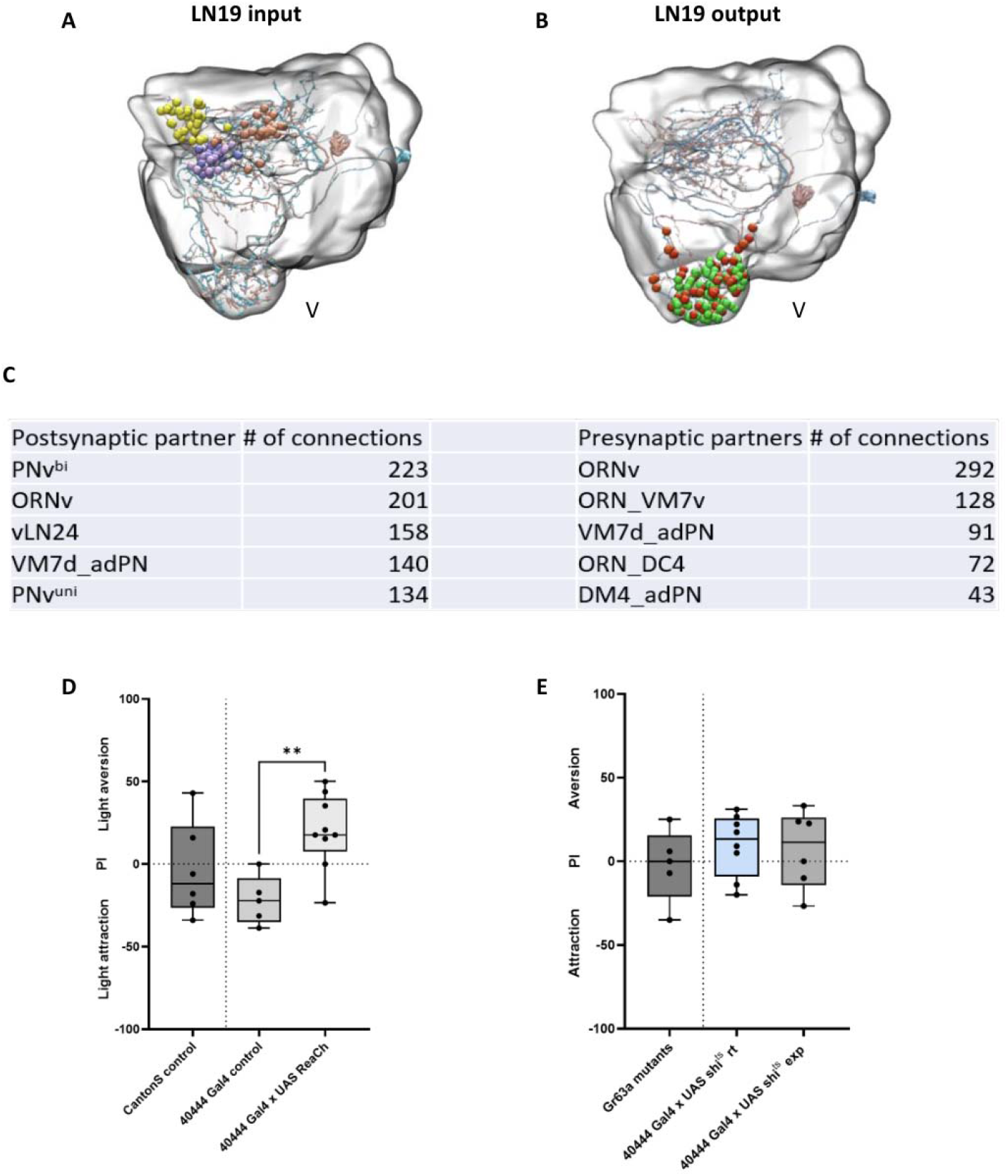
The “food”-odor-associated l2LN19. (A) LN19 receives anatomically identified input from DM4, DC4/DP1, and VM7, while (B) its output includes the V glomerulus. (C) Table illustrating the pre- and postsynaptic partners of LN19 according to the hemibrain connectome (neuPrint). (D) Optogenetic activation of LN19 using a specific driver line induced aversion in the behavioral choice assay. (E) Silencing of LN19 via shibirets did not alter innate CO_2_ avoidance under the tested conditions. “rt” indicates the room-temperature control for shibirets, while “exp” indicates the restrictive-temperature condition. These manipulations did not directly test the effect of adding food odor to CO_2_ stimulation.

Optogenetic activation of the two tested PNm populations downstream of LN23 induced opposite behavioral responses: While activity of the “CO_2_-only” class PNm1 results in strong aversion, the “multi-modal” PNm17 initiates behavioral attraction (Fig.6D). Furthermore, CO_2_-induced aversive behavior could be attenuated following shits-mediated silencing of PNm1 but was not changed by silencing PNm17 (Fig.6 E, F). Interestingly, FlyWire.ai (Dorkenwald et al. 2024; Eckstein et al. 2024; Schlegel et al. 2024; Zheng et al. 2018) predicted different neurotransmitter identities for the two PNm classes, with a GABAergic prediction for PNm1 and a cholinergic prediction for PNm17. Together with the behavioral effects of optogenetic activation, these data identify candidate nodes for opposing behavioral outputs; the proposed inhibitory mechanism is considered in the Discussion (Dolan et al. 2019).

### Connectivity in higher brain centers and lateral communication in the antennal lobe

As the glomerular PNs and the extra-glomerular LN23→PNm channels target adjacent central brain regions we were wondering if the two parallel CO_2_ pathways converge onto common downstream neurons. The hemibrain dataset (https://neuprint.janelia.org/; Plaza et al. 2022; Scheffer et al. 2020) identified three main types of glutamatergic LH neurons (FlyWire.ai; Dorkenwald et al. 2024; Eckstein et al. 2024; Schlegel et al. 2024; Zheng et al. 2018) as PNv or PNm targets: LHPD5c1 as the primary target of PNv^bi^ (Fig.6 G), LHPV9b1 downstream of PNm1 (Fig.6 H) and LHCENT4 as the major postsynaptic partner of PNv^uni^ (Fig.6 I). All 3 LH target neurons of the parallel CO_2_ pathways are directly interconnected (Fig. 8): While LHPV9b1 not only receives input from PNm1 but also from PNv^bi^, LHPD5c1, the main downstream target of PNv^bi^, is also targeted by LHCENT4, the primary postsynaptic partner of PNv^uni^. This core motif of CO_2_ perception receives additional input from other sensory modalities. LHPD5c1 gains input from “food-sensing” glomeruli (DM4, DC4/DP1, and VM7) via class-specific cholinergic PNs, providing anatomical evidence for convergence of CO_2_ and food-associated channels (Couto et al. 2005; Dumenil et al. 2025; Min et al. 2013). In addition, LHCENT4 is also a main target for l2PNVP4+VL1, mediating the perception of dryness and pyrrolidine-type odorants, suggesting a potential additional route for cross-modal interactions (Fig.8) (Li et al. 2022; Choi et al. 2022; Ni 2021). Interestingly, the convergence of the glomerular CO_2_ and food channels is not only observed for LH neurons downstream of AL relay pathways but seems to occur already at the level of AL glomeruli. Hemibrain describes with l2LN19 another class of polarized local interneuron (Fig.7A, B) with enriched postsynaptic sites at “food-sensing” glomeruli DM4, DC4/DP1, and VM7 (Fig.7A) as well as targeted output onto PNv^bi^ and PNv^uni^ in the V-glomerulus (Fig.7B), but merely with LN23 (Fig.7C) (https://neuprint.janelia.org/; Schlegel et al. 2021; Bates et al.2020). Optogenetic activation of LN19 led to an aversive response (Fig.7D), whereas LN19 silencing did not alter CO_2_ avoidance under the tested conditions (Fig.7E).

**Figure 8.**
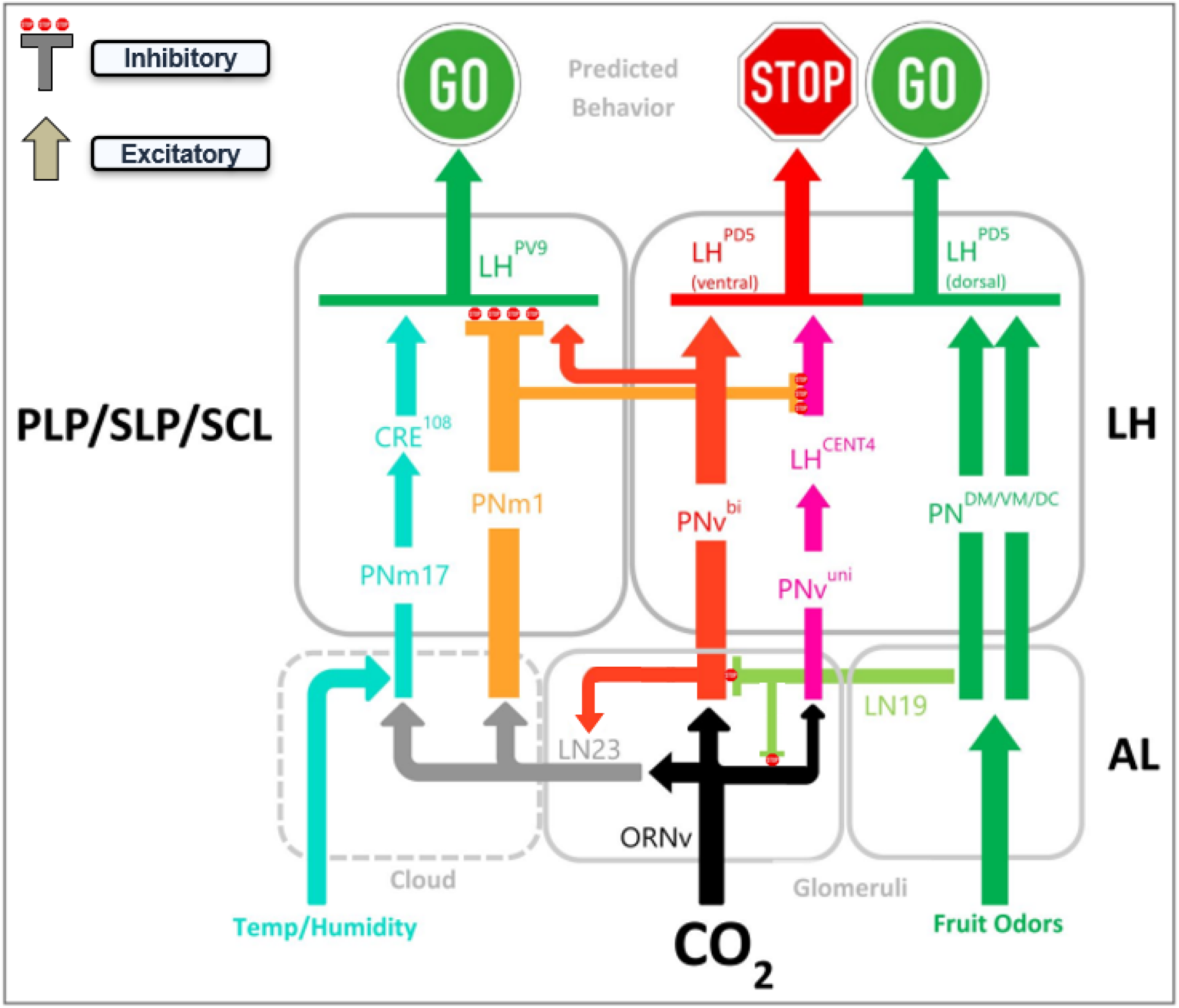
Working model of parallel channels for CO_2_ valence coding. The schematic distinguishes the canonical glomerular outputs PNv^bi^ and PNv^uni^ from the LN23-associated extraglomerular route to PNm1 and PNm17, and depicts potential convergence with food-odor and thermo-/hygrosensory channels. A PNv^bi^→LN23 connection is shown, consistent with the activity-associated sybGRASP signal observed in Figure 2; its functional strength and effect on LN23 activity were not directly measured. The GO/STOP symbols indicate hypothesized downstream behavioral effects rather than identified motor-command neurons. Inhibitory and excitatory signs, including the proposed PNm1-mediated suppression of attraction-related output, are model predictions informed partly by connectomic connectivity and neurotransmitter predictions; the complete context-dependent circuit computation was not experimentally established.

Figure 8 summarizes the anatomical pathways, neuronal manipulation results, and proposed higher-order interactions as a working circuit model; its hypothesized synaptic signs and behavioral balance are discussed below.

## Discussion

In *Drosophila*, the activation of a single class of CO_2_ receptive sensory neurons induces avoidance behavior, which is modified by multiple external and internal stimuli (Bräcker et al. 2013; Faucher et al. 2006; Dissegna et al. 2021; Suh et al. 2004; Turner and Ray, 2009). How does the *Drosophila* brain transform ORNv input into flexible behavioral responses towards CO_2_? Our findings extend the canonical glomerular model by identifying LN23 as an additional polarized relay through which ORNv information reaches a posterior extraglomerular region of the antennal lobe. This organization suggests that a single sensory channel can diverge into parallel routes before higher-order processing, potentially providing distinct sites for modulation. The comparatively small changes in the tested LN23 CaLexA and presynaptic-marker measures after prolonged CO_2_ exposure, relative to PNv^bi^, might reflect intrinsic cellular properties or differences in GABAergic modulation; these reporter measurements do not establish invariant LN23 firing. The prominent behavioral effect of LN23 manipulation, despite the more modest effects of manipulating individual canonical PNs, raises the question of how these routes are weighted and interact to generate avoidance. The predicted GABAergic identity of PNm1 (FlyWire.ai; Dorkenwald et al. 2024; Eckstein et al. 2024; Schlegel et al. 2024; Zheng et al. 2018) offers one possible downstream mechanism through inhibition of attraction-related signaling, but the relevant postsynaptic targets and physiological effect have not been directly tested.

An unresolved question is how the strong effect of LN23 silencing can coexist with the aversive contributions of the canonical PN pathways. The attraction observed after prolonged CO_2_ exposure in the activity-dependent LN23-silencing condition may reflect the balance of remaining pathways, but this experiment combines prolonged exposure with conditional silencing and does not establish that either PN class intrinsically mediates attraction. The increase in PNv^bi^ →LN23 sybGRASP following exposure suggests altered activity-associated contacts between the routes; it does not demonstrate compensatory signaling, increased LN23 firing, or preservation of aversion (Sachse et al. 2007). Although PNv^uni^ and PNv^bi^ are both described as cholinergic projection neurons (Tanaka et al. 2012; Lin et al. 2013; Bates et al. 2020; Schlegel et al. 2021), their different distributions of presynaptic markers suggest distinct opportunities for feed-forward transmission and intraglomerular modulation. Thus, LN23, PNv^uni^, and PNv^bi^ are likely to make complementary rather than interchangeable contributions to CO_2_ behavior, while the direction and strength of their interactions remain unresolved.

The neurotransmitter identity of LN23 remains unresolved, as the absence of detectable anti-GABA immunolabeling alone does not identify its transmitter (Supplementary Fig. 3A). One possibility is that LN23 signals via glutamate. In *Drosophila*, glutamate can have excitatory or inhibitory effects depending on the receptors expressed by postsynaptic neurons, including inhibitory glutamate-gated chloride channels (GluClα) characterized in the antennal lobe (Liu & Wilson, 2013). Thus, even if LN23 were glutamatergic, its functional effect on individual downstream partners could not be inferred from transmitter identity alone.

Interestingly, developmental analysis revealed a polarized growth pattern of LN23 that resembles projection neurons more than typical local interneurons (Jefferis et al. 2004; Komiyama & Luo 2006). LN23 originates from a larval interneuron located in the suboesophageal zone (SEZ), which undergoes remodeling into an adult olfactory interneuron during early pupal stages (Truman & Bate, 1988). We hypothesize that the secreted Wnt5 protein, a critical factor for initial projection neuron dendrite patterning, supports the targeting of LN23 processes to the posterior antennal lobe (AL) region. Like the genetically defined recognition mechanisms that guide glomerulus-specific ORN-PN connectivity, growing ORNv axons appear to target LN23 dendrites as synaptic partners. Since LN23 dendrites are organized into a defined glomerular input and extraglomerular output zone, we propose Flamingo-dependent expression on ingrowing ORN axons to prevent sensory input to LN23 outside the glomerular domain.

The divergence of CO_2_-induced ORN activation into parallel relay channels provides anatomical opportunities for convergence with additional sensory information. This includes not only potential convergence of non-olfactory cues onto the PNm17-associated branch downstream of LN23 (Frank et al. 2015; Li et al. 2022), but also the sampling of food-odor-sensitive glomeruli by the polarized interneuron LN19 for V-glomerulus input (Couto et al. 2005; Dumenil et al. 2025; Min et al. 2013). The convergence of different parallel CO_2_ channels onto a common set of lateral horn (LH) interneurons suggests a potential hub for motor action control (Dolan et al. 2019). The results presented in this study support a conceptual model in which LN23-associated signaling could suppress attraction-related output through a PNm1-linked pathway and be modulated by additional glomerular and extraglomerular inputs; direct functional testing of this network computation is still required. Further studies on the physiological architecture of the CO_2_ core circuit within and downstream of the LH may reveal deeper insights into the multimodal processing underlying flexible, context-dependent behaviors. Although chronic CO_2_ exposure reduced avoidance, the present experiments do not distinguish habituation from sensory adaptation, fatigue, or other prolonged-exposure effects. CaLexA cannot resolve fast responses or establish constant LN23 firing, and the lower baseline avoidance in the LN23 CaLexA genetic background (Figure 5B) warrants caution when assigning the entire shift to conditional silencing. The experiments also did not independently manipulate food-odor context, so the model does not establish food-dependent switching of CO_2_ valence.

Additionally, despite the conserved molecular receptor identity of CO_2_ sensory neurons among insects, species-specific behavioral responses to local CO_2_ concentrations are evident (Turner & Ray 2009). In many species, such as mosquitoes, elevated CO_2_ levels signal potential food sources and elicit attraction (Tauxe et al. 2013; Siju 2009; Ghaninia 2007). The identification of a dedicated CO_2_ aversion channel in D. melanogaster opens new avenues for comparative research into circuit motifs and offers insight into the neural basis of behavioral flexibility across insect evolution (Oram & Card 2022).

---

**Supplementary Figure 1.**
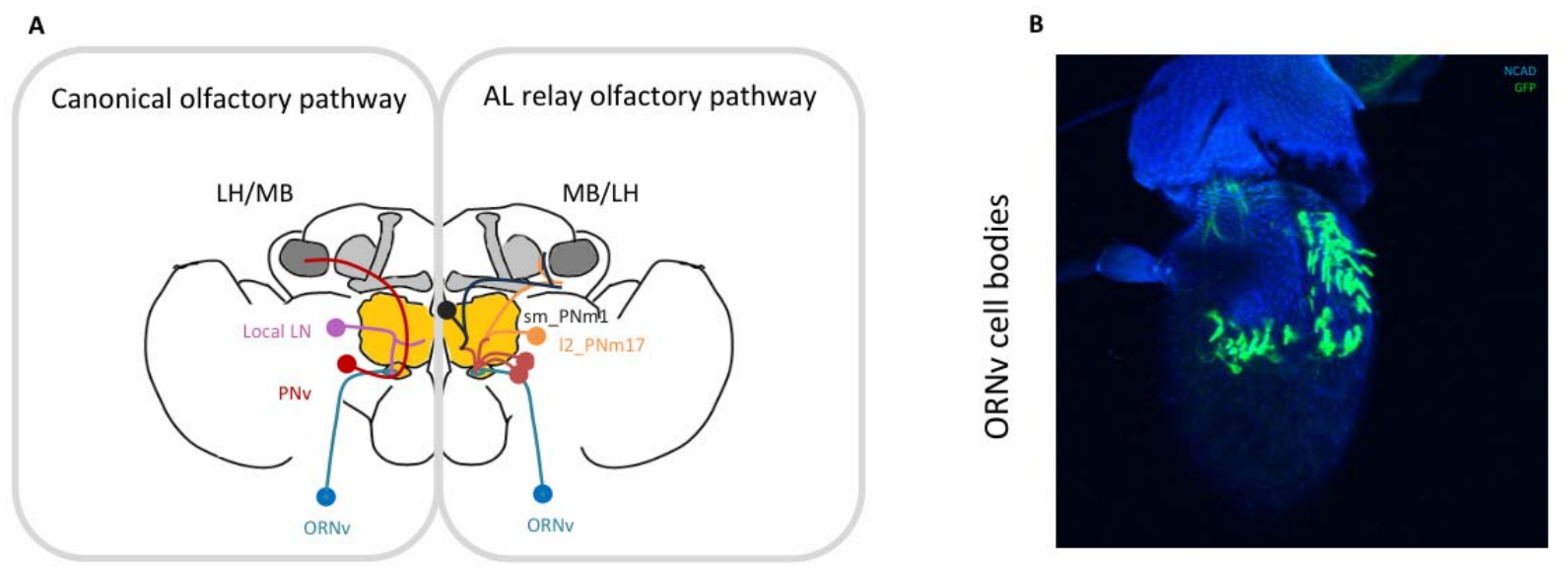
(A) Schematic of the canonical olfactory pathway (left) and the identified AL relay pathway (right). (B) CaLexA::GFP expression in vORN cell bodies on the third antennal segment.

**Supplementary Figure 3.**
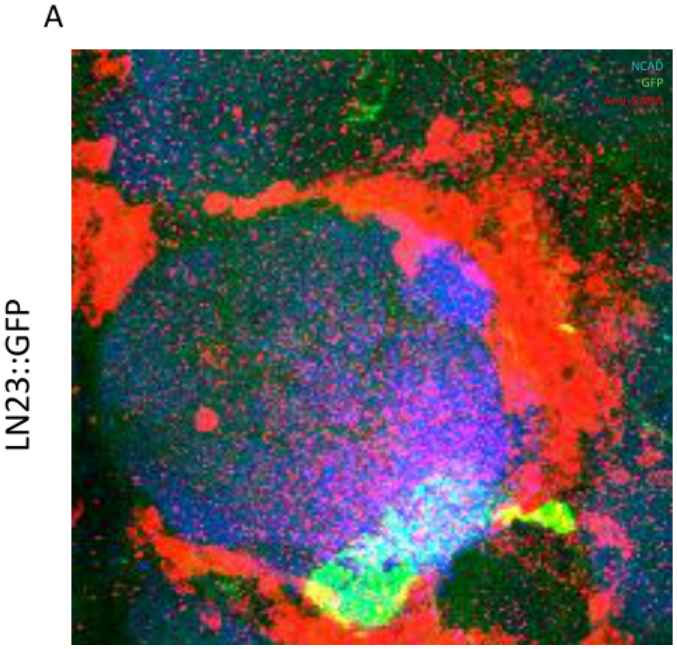
(A) CaLexA labeling of LN23 combined with anti-GABA immunostaining did not show detectable colocalization in the preparation examined. This negative result does not alone establish that LN23 lacks all inhibitory signaling.

**Supplementary Figure 4.**
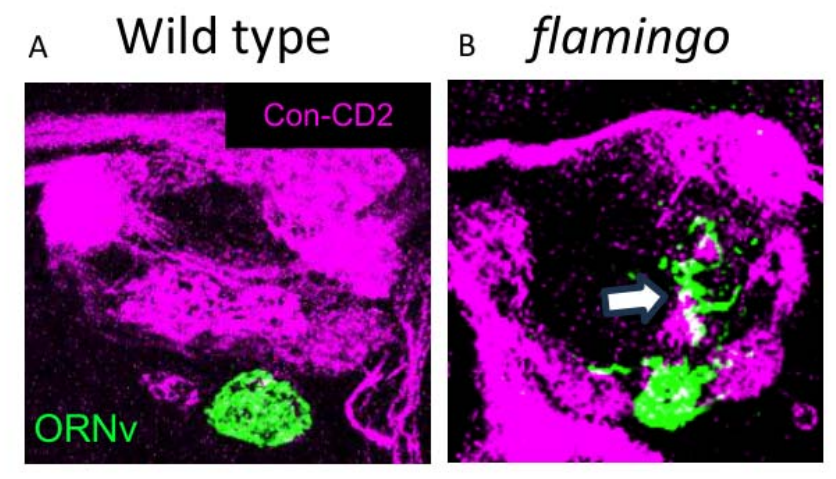
Loss of the cell adhesion molecule Flamingo in ORNv. In contrast to (A) wild-type axonal convergence of ORNv within the glomerular boundaries, (B) fmi-mutant ORNv axons overshoot the ventral glomerular domain and extend into the posterior AL (arrow).

